# Real-time dynamic single-molecule protein sequencing on an integrated semiconductor device

**DOI:** 10.1101/2022.01.04.475002

**Authors:** Brian D. Reed, Michael J. Meyer, Valentin Abramzon, Omer Ad, Pat Adcock, Faisal R. Ahmad, Gün Alppay, James A. Ball, James Beach, Dominique Belhachemi, Anthony Bellofiore, Michael Bellos, Juan Felipe Beltrán, Andrew Betts, Mohammad Wadud Bhuiya, Kristin Blacklock, Robert Boer, David Boisvert, Norman D. Brault, Aaron Buxbaum, Steve Caprio, Changhoon Choi, Thomas D. Christian, Robert Clancy, Joseph Clark, Thomas Connolly, Kathren Fink Croce, Richard Cullen, Mel Davey, Jack Davidson, Mohamed M. Elshenawy, Michael Ferrigno, Daniel Frier, Saketh Gudipati, Stephanie Hamill, Zhaoyu He, Sharath Hosali, Haidong Huang, Le Huang, Ali Kabiri, Gennadiy Kriger, Brittany Lathrop, An Li, Peter Lim, Stephen Liu, Feixiang Luo, Caixia Lv, Xiaoxiao Ma, Evan McCormack, Michele Millham, Roger Nani, Manjula Pandey, John Parillo, Gayatri Patel, Douglas H. Pike, Kyle Preston, Adeline Pichard-Kostuch, Kyle Rearick, Todd Rearick, Marco Ribezzi-Crivellari, Gerard Schmid, Jonathan Schultz, Xinghua Shi, Badri Singh, Nikita Srivastava, Shannon F. Stewman, T.R. Thurston, Philip Trioli, Jennifer Tullman, Xin Wang, Yen-Chih Wang, Eric A. G. Webster, Zhizhuo Zhang, Jorge Zuniga, Smita S. Patel, Andrew D. Griffiths, Antoine M. van Oijen, Michael McKenna, Matthew D. Dyer, Jonathan M. Rothberg

**Affiliations:** Quantum-Si, Inc., 530 Old Whitfield Street, Guilford, Connecticut 06437, USA; Laboratoire de Biochimie, ESPCI Paris, Université PSL, CNRS UMR 8231, 10 Rue Vauquelin, Paris, France; Department of Biochemistry and Molecular Biology, Rutgers University, 683 Hoes Lane, Piscataway, New Jersey 08854, USA; University of Wollongong, Wollongong, NSW 2552, Australia

## Abstract

Proteins are the main structural and functional components of cells, and their dynamic regulation and post-translational modifications (PTMs) underlie cellular phenotypes. Next-generation DNA sequencing technologies have revolutionized our understanding of heredity and gene regulation, but the complex and dynamic states of cells are not fully captured by the genome and transcriptome. Sensitive measurements of the proteome are needed to fully understand biological processes and changes to the proteome that occur in disease states. Studies of the proteome would benefit greatly from methods to directly sequence and digitally quantify proteins and detect PTMs with single-molecule sensitivity and precision. However current methods for studying the proteome lag behind DNA sequencing in throughput, sensitivity, and accessibility due to the complexity and dynamic range of the proteome, the chemical properties of proteins, and the inability to amplify proteins. Here, we demonstrate single-molecule protein sequencing on a compact benchtop instrument using a dynamic sequencing by stepwise degradation approach in which single surface-immobilized peptide molecules are probed in real-time by a mixture of dye-labeled N-terminal amino acid recognizers and simultaneously cleaved by aminopeptidases. By measuring fluorescence intensity, lifetime, and binding kinetics of recognizers on an integrated semiconductor chip we are able to annotate amino acids and identify the peptide sequence. We describe the expansion of the number of recognizable amino acids and demonstrate the kinetic principles that allow individual recognizers to identify multiple amino acids in a highly information-rich manner that is sensitive to adjacent residues. Furthermore, we demonstrate that our method is compatible with both synthetic and natural peptides, and capable of detecting single amino acid changes and PTMs. We anticipate that with further development our protein sequencing method will offer a sensitive, scalable, and accessible platform for studies of the proteome.

## Main Manuscript

Measurements of the proteome provide deep and valuable insight into biological processes. However, the complex nature of the proteome and the chemical properties of proteins present several fundamental challenges to achieving sensitivity, throughput, cost, and adoption on par with DNA sequencing technologies^1,2^. These challenges include the large number of different proteins per cell (>10,000) and perhaps 10-fold larger number of proteoforms^3^; the very wide dynamic range of protein abundance in cells and biological fluids^4,5^ and lack of correlation with transcript levels^6^; the costs and high detection limits of current mass spectrometry methods^2^; and the inability to copy or amplify proteins.

Here, we present a single-molecule protein sequencing approach and integrated system for massively-parallel proteomic studies. We immobilize peptides in nanoscale reaction chambers on a semiconductor chip and detect N-terminal amino acids (NAAs) with dye-labeled NAA recognizers in real-time. Aminopeptidases sequentially remove individual NAAs to expose subsequent amino acids for recognition, eliminating the need for complex chemistry and fluidics (Fig. 1). We built a benchtop device with a 532 nm pulsed laser source for fluorescence excitation and electronics for signal processing (Extended Data Fig. 1a). Our semiconductor chip uses fluorescence intensity and lifetime, rather than emission wavelength, for discrimination of dye labels. Our recognizers detect one or more types of NAAs and provide information for peptide identification based on the temporal order of NAA recognition and the kinetics of on-off binding.

**Figure 1.**
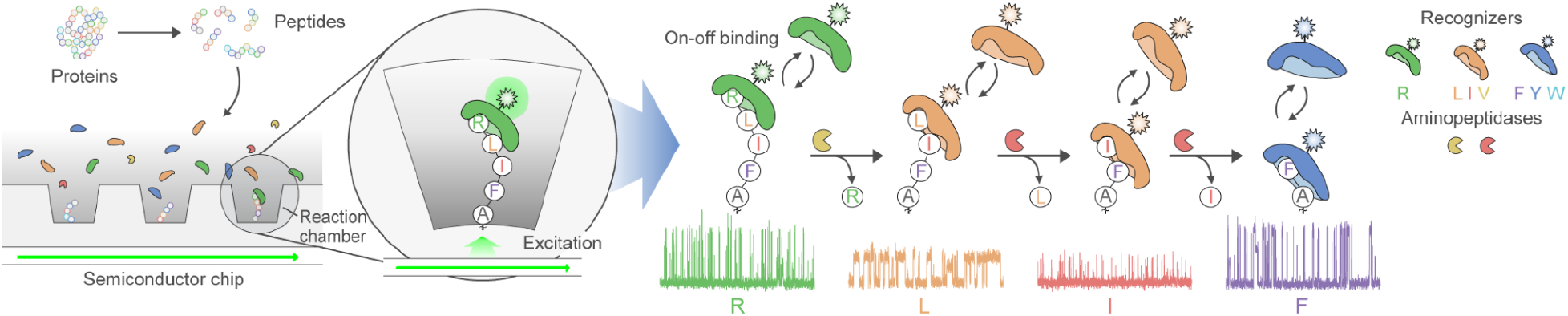
Overview of real-time dynamic protein sequencing. Protein samples are digested into peptide fragments, immobilized in nanoscale reaction chambers, and incubated with a mixture of freely-diffusing NAA recognizers and aminopeptidases that carry out the sequencing process. The labeled recognizers bind on and off to the peptide when one of their cognate NAAs is exposed at the N-terminus, thereby producing characteristic pulsing patterns. The NAA is cleaved by an aminopeptidase, exposing the next amino acid for recognition. The temporal order of NAA recognition and the kinetics of binding enable peptide identification and are sensitive to features that modulate binding kinetics, such as PTMs.

We apply this sequencing approach to peptides of diverse sequence composition and length. We also demonstrate the ability to resolve single amino acid substitutions and PTMs based on the kinetic signatures of recognizers, and to identify peptides in mixtures. Finally, we demonstrate sequencing of peptides derived from digestion of recombinant human proteins. The core sequencing technology we present here provides the basis for a sensitive and scalable platform capable of tackling the challenges of studying the proteome.

### A CMOS chip and integrated system for single-molecule measurements

We used complementary metal-oxide-semiconductor (CMOS) fabrication technology to build a custom time-domain-sensitive semiconductor chip with nanosecond precision, containing fully-integrated components for single-molecule detection, including photosensors, optical waveguide circuitry, and reaction chambers for biomolecule immobilization (Fig. 1). We achieve observation volumes less than 5 attoliters through evanescent illumination at reaction chamber bottoms from the nearby waveguide, enabling sensitive single-molecule detection in the context of high freely-diffusing dye concentrations (>1 μM).

The semiconductor chip utilizes a filterless system that excludes excitation light on the basis of photon arrival time, achieving greater than 10,000-fold attenuation of incident excitation light. Elimination of an integrated optical filter layer increases the efficiency of fluorescence collection and enables scalable manufacturing of the chip. To discriminate fluorescent dye labels attached to NAA recognizers by fluorescence lifetime and intensity, the chip rapidly alternates between early and late signal collection windows associated with each laser pulse, thereby collecting different portions of the exponential fluorescence lifetime decay curve. The relative signal in these collection windows (termed ‘bin ratio’) provides a reliable indication of fluorescence lifetime (Extended Data Fig. 1b-f, Methods).

### Ordered recognition and cleavage of NAAs on single peptide molecules in real-time

For NAA binding proteins to function as recognizers, the recognizer-peptide complex must remain bound long enough (typically >120 ms on average) to generate detectable single-molecule binding events. We first focused on proteins from the N-end rule adapter family ClpS that naturally bind to N-terminal phenylalanine, tyrosine, and tryptophan^7–9^. Using PS610, a recognizer we derived from ClpS2 from *Agrobacterium tumefaciens*, we established that this recognizer binds detectably to immobilized peptides with these NAAs. Importantly, we also determined that the kinetics of binding differ for each NAA. To demonstrate these properties, we incubated immobilized peptides containing the initial N-terminal sequences FAA, YAA, or WAA on separate chips with PS610 and collected data for 10 hours (Methods). We observed NAA recognition by PS610, characterized by continuous on-off binding during the incubation period, with distinct pulse duration (PD) for each peptide (Fig. 2a). Median PDs were 2.49, 0.73, and 0.31 s for FAA, YAA, and WAA, respectively. These values reflect differences in binding affinity driven by different dissociation rates for each type of protein-NAA interaction7 (Extended Data Fig. 2a,b).

**Figure 2.**
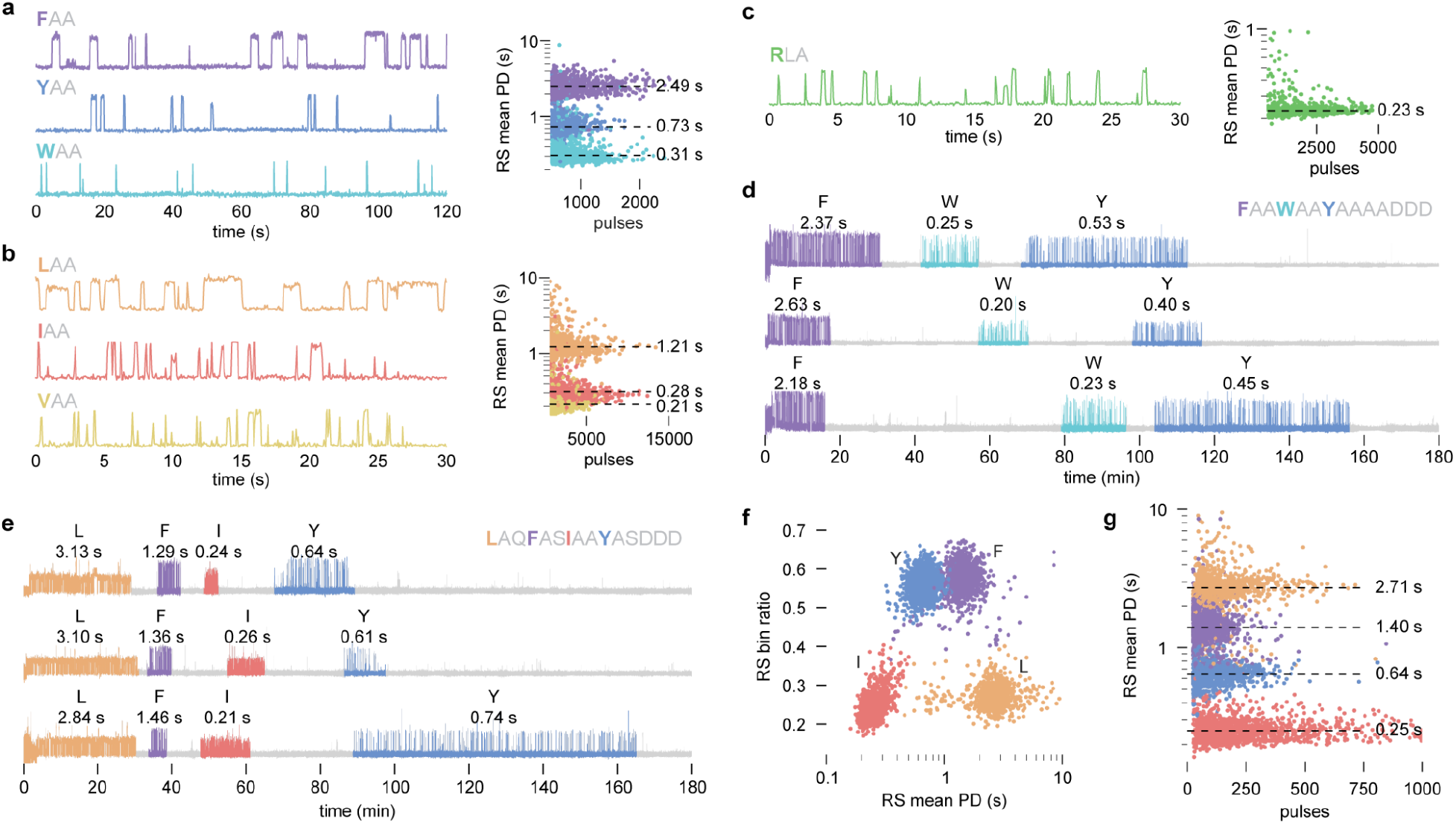
NAA recognition and dynamic sequencing. **a**-**c**, Example traces demonstrating single-molecule N-terminal recognition by PS610 (**a**), PS961 (**b**), and PS691 (**c**). Scatter plots of the number of pulses per RS vs RS mean PD are displayed for each peptide in **a**-**c**, with median PD indicated. **d**, Example traces from dynamic sequencing of the synthetic peptide FAAWAAYAADDD. Median PD is indicated above each RS. **e-g**, Dynamic sequencing of the synthetic peptide LAQFASIAAYASDDD using PS610 and PS961. **e**, Example traces. **f**, Scatter plot of RS mean PD vs bin ratio illustrating discrimination of recognizers by bin ratio and NAAs by pulse duration. **g**, Scatter plot of the number of pulses per RS vs RS mean PD, grouped by the amino acid label assigned to the RS.

To expand the set of recognizable NAAs, we further investigated N-end rule pathway proteins as a source of additional recognizers. In a comprehensive screen of diverse ClpS family proteins, we discovered a group of ClpS proteins from the bacterial phylum Planctomycetes with native binding to N-terminal leucine, isoleucine, and valine. We applied directed evolution techniques to generate a Planctomycetes ClpS variant—PS961—with sub-micromolar affinity to N-terminal leucine, isoleucine, and valine, and demonstrated recognition of these NAAs (Fig. 2b). The median PD of binding to peptides with N-terminal LAA, IAA, and VAA was 1.21, 0.28, and 0.21 s, respectively, in agreement with bulk characterization (Extended Data Fig. 2c).

In a separate screen, we investigated a diverse set of UBR-box domains from the UBR family of ubiquitin ligases that natively bind N-terminal arginine, lysine, and histidine^10^. The UBR-box domain from the yeast *Kluyveromyces lactis* UBR1 protein exhibited the highest affinity for N-terminal arginine, and we used this protein to generate an arginine recognizer, PS691. PS691 recognized arginine in a peptide with N-terminal RLA with a median PD of 0.23 s (Fig. 2c). Lower affinity binding to N-terminal lysine and histidine (Extended Data Fig. 2d,e) was insufficient for single-molecule detection.

To demonstrate that amino acids in a single peptide molecule can be sequentially exposed by aminopeptidases and recognized in real-time with distinguishable kinetics, we incubated an immobilized peptide containing the initial sequence FAAWAAYAA with PS610 for 15 minutes, followed by addition of PhTET3, an aminopeptidase from *Pyrococcus horikoshii*^11^. The collected traces consisted of regions of distinct pulsing, which we refer to as recognition segments (RSs), separated by regions lacking recognition pulsing (non-recognition segments, NRSs). We developed analysis software to automatically identify pulsing regions and transition points within traces (Methods). Traces began with recognition of phenylalanine with median PD of 2.36 s (Fig. 2d), in agreement with the PD observed for FAA in recognition-only assays. This pattern terminated after aminopeptidase addition (on average 11 min after addition), and was followed by the ordered appearance of two RSs with median PD of 0.25 s and 0.49 s (Fig. 2d), corresponding to the short and medium PDs obtained in our YAA and WAA recognition-only assays. Thus, the introduction of aminopeptidase activity to the reaction resulted in the sequential appearance of discrete RSs with the expected kinetic properties in the correct order.

To demonstrate dynamic sequencing with two NAA recognizers, we labeled PS610 and PS961 with the distinguishable dyes atto-Rho6G and Cy3, respectively, and exposed an immobilized peptide of sequence LAQFASIAAYASDDD to a solution containing both recognizers. After 15 minutes, we added two *P. horikoshii* aminopeptidases with combined activity covering all 20 amino acids—PhTET2 and PhTET3^11,12^. The collected traces displayed discrete segments of pulsing alternating between PS961 and PS610 according to the order of recognizable amino acids in the peptide sequence (Fig. 2e). The average bin ratio and average PD associated with each RS readily distinguished the two dye labels and four types of recognized NAAs (Fig. 2f). Median PDs were 2.71, 1.40, 0.25, and 0.64 s for N-terminal LAQ, FAS, IAA, and YAS, respectively (Fig. 2g).

Previous studies have shown that NAA-bound ClpS and UBR proteins also make contacts with the residues at position 2 (P2) and position 3 (P3) from the N-terminus that influence binding affinity^9,13,14^. These influences are reflected in the modulation of PD depending on the downstream P2 and P3 residues, as we observed above for LAA (1.21 s) compared to LAQ (2.70 s). We find that these influences on PD vary within informatically advantageous ranges and can be determined empirically or approximated in silico to model peptide sequencing behavior a priori (Extended Data Fig. 2f-h). A powerful feature of this recognition behavior in regard to peptide identification is that each RS contains information about potential downstream P2 and P3 residues or PTMs, whether or not these positions are the targets of an NAA recognizer.

### Principles of dynamic protein sequencing illustrated with model peptides

To evaluate the kinetic principles of our dynamic sequencing method when applied to diverse sequences, we first characterized the synthetic peptide DQQRLIFAG, corresponding to a segment of human ubiquitin (Fig. 3a-e). We performed sequencing reactions using a combination of three differentially-labeled recognizers—PS610, PS961, and PS691—and two aminopeptidases—PhTET2 and PhTET3 (Methods). The example trace in Figure 3a starts with an NRS that corresponds to the time interval during which residues in the initial DQQ motif are present at the N-terminus. The first RS starts at 120 min, upon exposure of N-terminal arginine to recognition by PS691. Subsequent cleavage events sequentially expose N-terminal leucine, isoleucine, and phenylalanine to their corresponding recognizers, with fast transitions (average <10 s) from one RS to the next. This overall pattern is replicated across many instances of sequencing of the same peptide, with similar PD statistics across traces, as each peptide molecule follows the same reaction pathway over the course of the sequencing run (Fig. 3b-c). Due to the stochastic timing of cleavage events, each trace displays distinct start times and durations for each RS (Fig. 3c).

**Figure 3.**
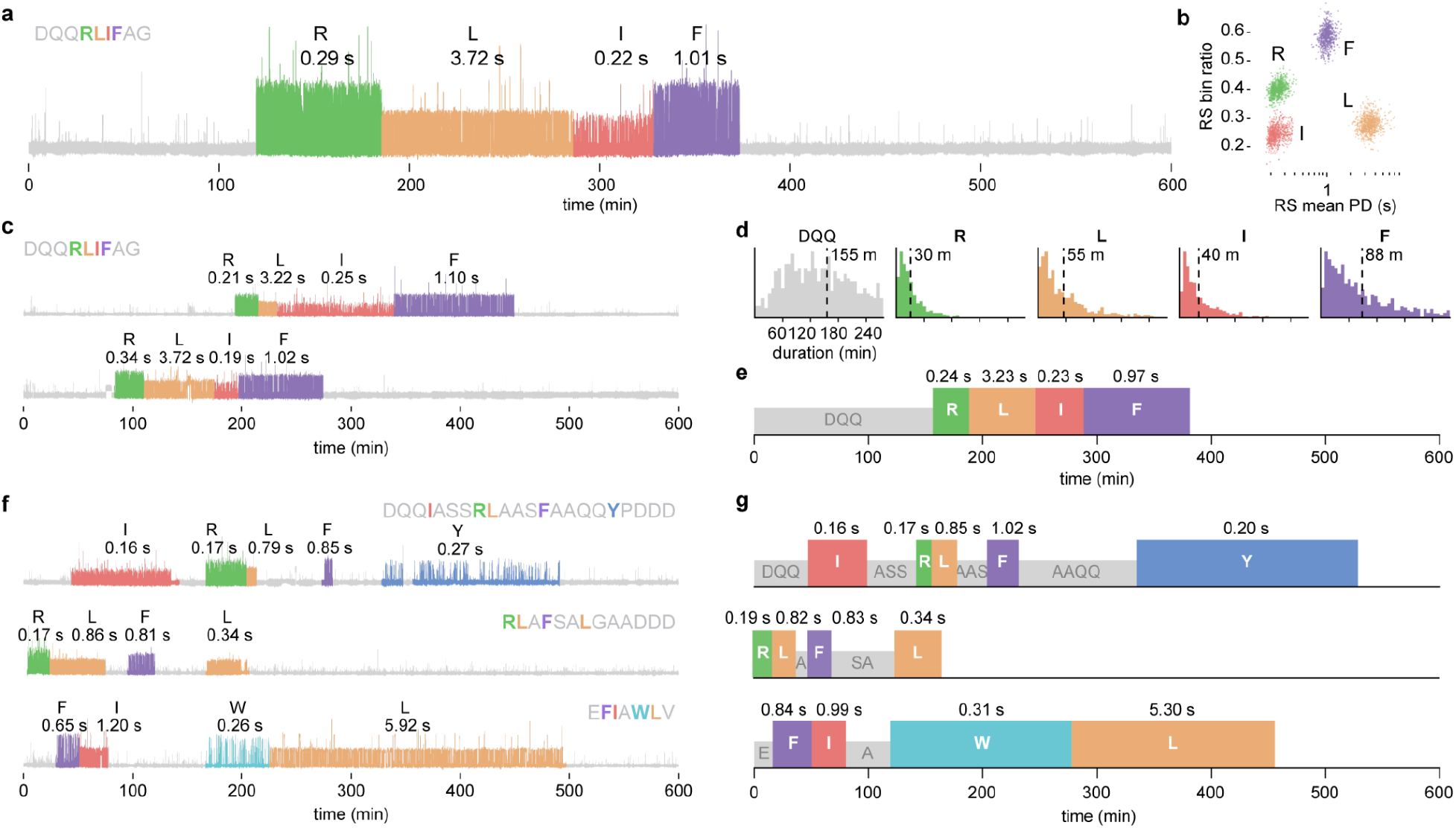
Dynamic sequencing of diverse peptides with high-precision kinetic outputs. **a-e**, Dynamic sequencing of the peptide DQQRLIFAG. **a**, Example trace. **b**, Scatter plot of RS mean PD vs bin ratio. **c**, Additional example traces of dynamic sequencing of DQQRLIFAG peptide. **d**, Distributions of the duration of each RS and NRS acquired during sequencing, with mean durations indicated. **e**, Kinetic signature plots summarizing the characteristic sequencing behavior of DQQRLIFAG peptide. **f-g**, Dynamic sequencing of the synthetic peptides DQQIASSRLAASFAAQQYPDDD (top), RLAFSALGAADDD (middle), and EFIAWLV (bottom). **f**, Example traces for each peptide. **g**, Corresponding kinetic signature plots.

This approach reports the binding kinetics at each recognizable amino acid position and the kinetics of aminopeptidase cleavage along the peptide sequence. High-precision kinetic information on binding is obtained from a single trace, since each RS typically contains tens to hundreds of on-off binding events, resulting in a distribution of PD and interpulse duration (IPD) measurements that can be analyzed statistically. The repetitive probing of each NAA also provides accurate recognizer calling, since calls are not based on the error-prone detection of a single event associated with one fluorophore molecule (Extended Data Fig. 1f). Recognizer on-rate and concentration governs IPD for each RS; higher recognizer concentrations result in shorter average IPDs and faster rates of pulsing (Extended Data Fig. 3a,b). Higher recognizer concentrations, however, increase the fluorescence background from freely diffusing recognizers, resulting in lower pulse signal-to-noise, and can compete with aminopeptidases for N-terminal access. In practice, IPDs in the range of approximately 2 to 10 s provide a favorable balance among these factors.

The distribution of RS durations across an ensemble of replicate traces defines the rate of cleavage of each recognizable NAA. For DQQRLIFAG peptide, we observed average cleavage times of 30, 55, 40, and 88 min for N-terminal arginine, leucine, isoleucine, and phenylalanine, respectively, with approximate single-exponential decay statistics for each position (Fig. 3d, Extended Data Fig. 3c). The distribution of NRS durations reports the cleavage rate of a run of one or more non-recognized NAAs. The average NRS duration for the initial DQQ motif was 155 min (Fig. 3d). Average cleavage rates are a key parameter and are controlled by the aminopeptidase concentration in the assay (Extended Data Fig. 3d,e). Given the exponential behavior, we target average RS durations of 10 to 40 min to provide sufficient time for pulsing data collection, avoid missed RSs due to rapid cleavage, and minimize excessively long RS durations. We found it helpful to visualize the sequencing profiles of peptides as kinetic signature plots—simplified trace-like representations of the time course of complete peptide sequencing containing the median PD for each RS, and the average duration of each RS and NRS (Fig. 3e). These highly characteristic features provide a wealth of sequence-dependent information for mapping traces from peptides to their proteins of origin.

To demonstrate that this core methodology and its kinetic principles apply to a wide range of peptide sequences, we sequenced the synthetic peptides DQQIASSRLAASFAAQQYPDDD, RLAFSALGAADDD, and EFIAWLV—a segment of human Glucagon-like peptide-1 (GLP-1)—under the same sequencing conditions used for DQQRLIFAG (Fig. 3f). Each peptide generated a characteristic kinetic signature in accordance with its sequence (Fig. 3g). We obtained readouts as far as position 18 (the furthest recognizable amino acid) in the peptide DQQIASSRLAASFAAQQYPDDD, illustrating that the method is compatible with long peptides and capable of deep access to sequence information in peptides.

### Distinct kinetic signatures from single amino acid changes and PTMs

To illustrate how the kinetic parameters acquired from sequencing are sensitive to changes in sequence composition, we performed sequencing with a set of three peptides that differ only at the third position—RLAFAYPDDD, RLIFAYPDDD, and RLVFAYPDDD (Fig. 4a). Each type of amino acid at this position had a distinct effect on the PD acquired during recognition of N-terminal leucine by PS961. We observed median PDs of 1.29 s, 2.22 s, and 4.21 s for LAF, LIF, and LVF, respectively (Fig. 4b). Importantly, these kinetic effects do not depend on whether the third residue is recognizable as an NAA, illustrating that the kinetic signatures of peptides contain information on sequence composition for both recognized NAAs and adjacent residues. In addition to differences in PD for leucine, each peptide displayed a characteristic RS or NRS in the interval between leucine and phenylalanine recognition (Fig. 4a, Extended Data Fig. 4a). These results demonstrate the sensitivity of the sequencing readout to single point substitutions.

**Figure 4.**
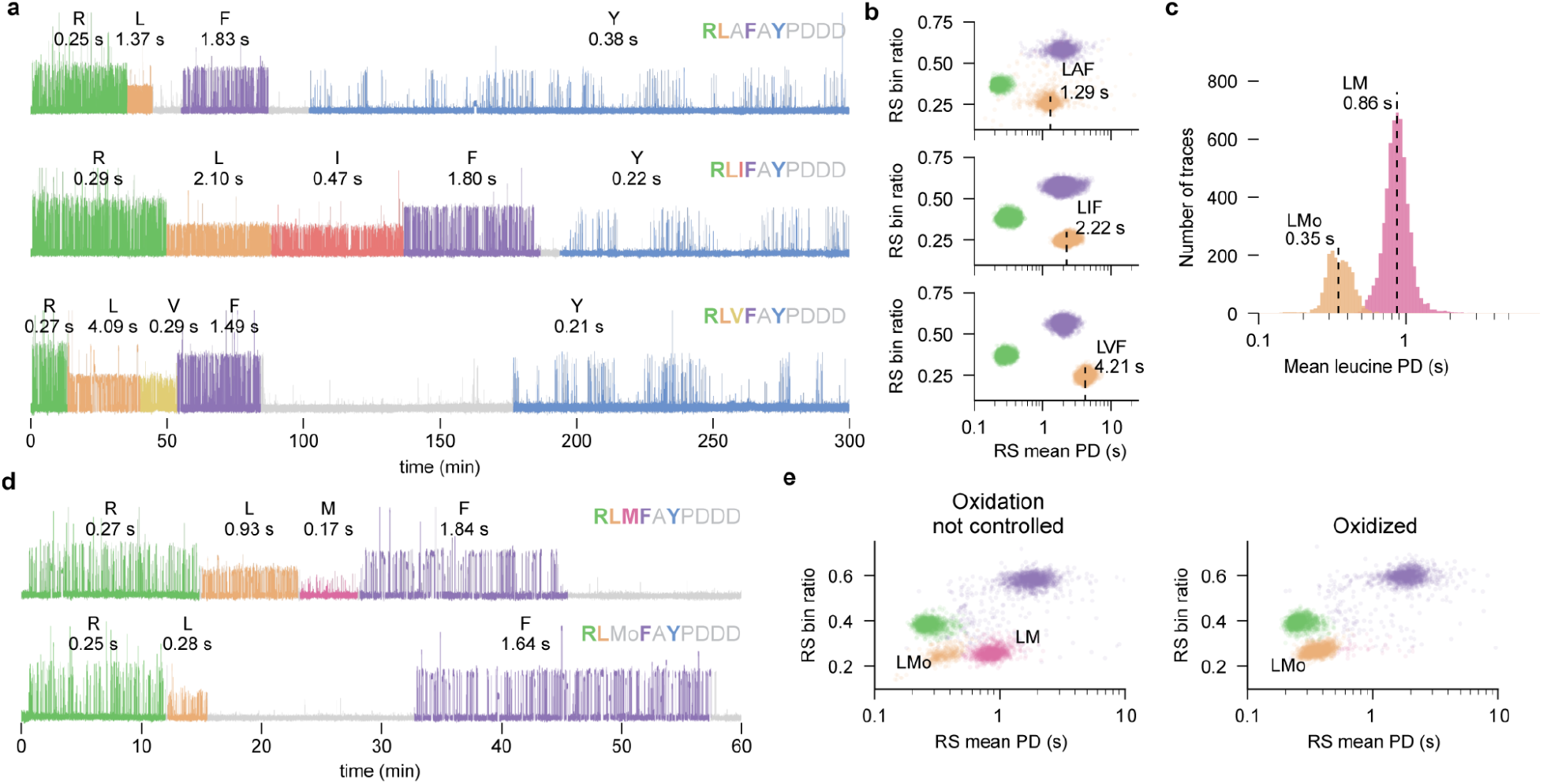
Detection of single amino acid changes and PTMs. **a-b**, Dynamic sequencing of synthetic peptides that differ by a single amino acid: RLAFAYPDDD (top), RLIFAYPDDD (middle), RLVFAYPDDD (bottom). **a**, Example traces. **b**, Scatter plots of RS mean PD vs bin ratio. **c-d**, Detection of oxidized methionine using the peptide RLMFAYPDDD. **c**, Distributions of mean PD for leucine; labels indicate populations with leucine followed by methionine (LM) or methionine sulfoxide (LMo). **d**, Example traces in which methionine is recognized by PS961 and leucine exhibits long PD (top), or in which methionine is not recognized due to oxidation and leucine exhibits short PD (bottom). **e**, Scatter plots of RS mean PD vs bin ratio for runs in which oxidation was not controlled (left) or in which methionine was fully oxidized (right).

Since the aminoacyl-proline bond of the YP motif in peptides such as RLIFAYPDDD cannot be cleaved by the PhTET aminopeptidases^11,12^, observation of YP pulsing at the end of a trace ensures that cleavage has progressed completely from the first to last recognizable amino acid. The sequencing output from RLIFAYPDDD, therefore, provided a convenient dataset for examining biochemical sources of non-ideal behavior that could lead to errors in peptide identification. The main sources of incomplete information in traces were deletions of expected RSs due to rapid sequential cleavage (Extended Data Fig. 4b) and early termination of reads resulting from photodamage or surface detachment (Extended Data Fig. 4c).

In addition to changes in amino acid sequence composition, sequencing readouts are sensitive to changes due to PTMs. As an example, we examined methionine oxidation. The thioether moiety of the methionine side chain is susceptible to oxidation during peptide synthesis and sequencing. We determined that PS961 binds a peptide with N-terminal methionine with a *K*_D_ of 947 ± 47 nM (Extended Data Fig. 4d) and hypothesized that oxidation, resulting in a polar methionine sulfoxide side chain, would eliminate binding and reduce NAA binding affinity when located at P2. We determined computationally that methionine sulfoxide is highly unfavorable in the PS961 NAA binding pocket and that non-polar residues are preferred at P2 (Extended Data Figs. 4e, 2h). We sequenced the synthetic peptide RLMFAYPDDD and observed two populations of traces with distinct kinetic signatures—a first population containing leucine recognition with median PD of 0.86 s, and a second population with median PD of 0.35 s (Fig. 4c). Traces from the first population also displayed methionine recognition with short PD in the time interval between leucine and phenylalanine recognition (Fig. 4d). Methionine recognition was absent in traces from the second population (Fig. 4d), indicating that the methionine side chain in these peptides was not capable of recognition by PS961. When we fully oxidized methionine by preincubation with hydrogen peroxide (Methods), we observed elimination of both methionine recognition and of the leucine recognition cluster with long median PD, as expected (Fig. 4e). These results demonstrate the capability for extremely sensitive detection of PTMs due to their kinetic effects on recognition.

### Sequencing peptide mixtures and mapping peptides derived from human proteins

Proteomics applications require identification of peptides in mixtures derived from biological sources. To extend our results to peptide mixtures and biologically-derived peptides, we performed two experiments. First, we mixed DQQRLIFAG and RLAFSALGAADDD peptides, immobilized them on the same chip, and performed a sequencing run. Data analysis (Methods) identified two populations of traces corresponding to each peptide, with kinetic signatures in close agreement with those identified in runs with individual peptides (Fig. 5a, Extended Data Fig. 4f). Second, to demonstrate that our method extends to biologically derived peptides, we performed sequencing runs with peptide libraries generated using a simple workflow from recombinant human ubiquitin (76 amino acids) and GLP-1 (37 amino acids) proteins digested with AspN/LysC and trypsin, respectively (Methods). For both libraries, data analysis readily identified traces matching the expected recognition pattern for the protease cleavage products DQQRLIFAGK and EFIAWLVK for ubiquitin and GLP-1, respectively, and produced kinetic signatures in agreement with synthetic versions of these peptides (Fig. 5b, Extended Data Fig. 4g). We identified matches to the kinetic signature of the ubiquitin peptide DQQRLIFAGK across the human proteome, taking advantage of simple sequence constraints provided by kinetic information (Methods). We found only one protein other than ubiquitin containing a peptide that could potentially match this signature (Fig. 5c); thus even short signatures can exhibit proteome abundance of less than one in 10^4^ proteins. These results illustrate the potential of the full kinetic output from sequencing to enable digital mapping of peptides to their proteins of origin.

**Figure 5.**
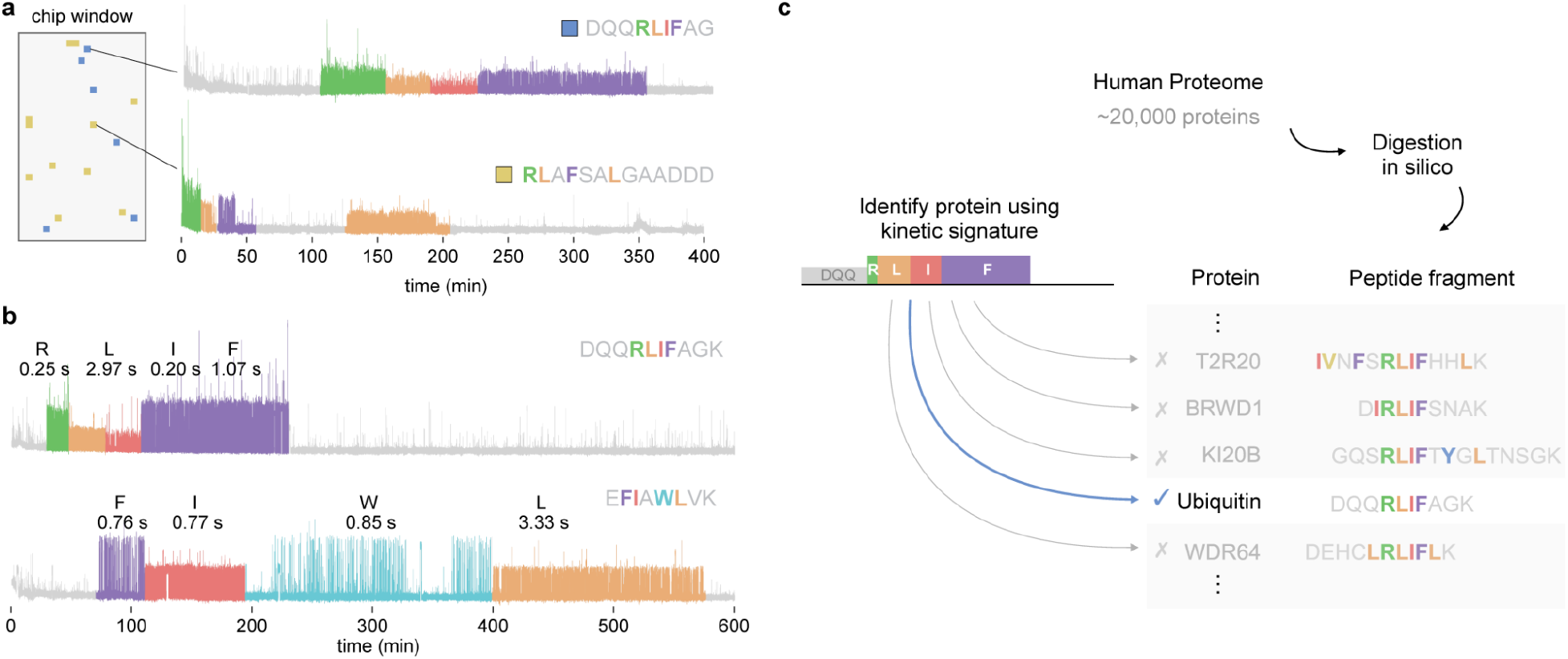
Discrimination of peptides in mixtures and mapping peptides to the human proteome. **a**, Example traces from sequencing a mixture of the peptides DQQRLIFAG and RLAFSALGAADDD on the same chip; the chip window indicates the location of reaction chambers producing a sequencing readout for each peptide. **b**, Example traces from the dynamic sequencing of two peptides, DQQRLIFAGK (top) and EFIAWLVK (bottom), isolated from the recombinant human proteins ubiquitin and GLP-1, respectively. **c**, Diagram illustrating identification of the protein ubiquitin as a match to the kinetic signature from DQQRLIFAGK peptide in an in silico digest of the human proteome based on kinetic information.

## Discussion

We have demonstrated a robust methodology capable of sequencing peptides of diverse sequence and length based on massively-parallel fluorescence-based single-molecule detection. The information-rich sequencing output of our assay, comprising highly characteristic kinetic signatures, including influences from downstream residues, permits the sensitive detection of single amino acid substitutions and PTMs.

Importantly, we demonstrated sequencing of peptides derived from digestion of recombinant human proteins using a simple workflow. Our simple, real-time dynamic approach differs markedly from other recently described single-molecule approaches that rely on complex, iterative methods involving stepwise Edman chemistry or hundreds of cycles of epitope probing^15–17^; nanopore approaches offer the potential for real-time readouts and simplicity, but face substantial challenges related to the size and biophysical complexity of polypeptides^18–20^.

Our sequencing technology is readily expanded in its capabilities, and there are multiple areas for improvement. Expansion of the number of recognizable amino acids can be achieved through continued directed evolution and engineering of recognizers. Amino acids that are currently recognizable at the N-terminal position comprise approximately 36.5% of the amino acid sequence content of the human proteome. Partial sequence information is sufficient for most proteomics applications, which rely on mapping to pre-defined sets of candidate proteins^21^, and the vast majority of human proteins contain at least one peptide suitable for mapping with current recognizers; for example, 96.7% of human proteins contain one or more peptides (average 8.6 peptides) of ten amino acids or longer with four or more recognizable amino acids using an AspN/LysC digest. Expanded amino acid coverage will result in kinetic signatures with increased information content, thereby increasing the number of informative peptides per protein and the complexity of peptide mixtures that can be sequenced simultaneously. Recognizers for new amino acids or PTMs can be evolved from current recognizers or identified in screens of other scaffolds, such as other types of NAA- or PTM-binding proteins or aptamers. Extension to direct N-terminal detection of all 20 natural amino acids and multiple PTMs is feasible for sequencing without reference to a protein database. Aminopeptidases can be engineered to optimize cleavage rates. We envision that the dynamic range of samples and the applications most suitable for the system will tend to scale with the number of reaction chambers on the chip, and that compression of dynamic range will be necessary for certain applications. We anticipate that future developments of the platform will increase the accessibility of proteomics studies and enable new discoveries in biological and clinical research.

## Acknowledgements

The authors would like to thank Emily Chen for helpful discussions in preparing the manuscript.

## Author Contributions

All of the authors contributed to the design, execution, and/or analysis of experiments related to the device, chip, and/or chemistry described in this manuscript. B.D.R. wrote the manuscript with input from co-authors.

## Data Availability

The datasets generated and/or analyzed during the current study are not publicly available as they are property of Quantum-Si, Inc. Please contact the corresponding author, M.D.D., if you are interested in accessing the data.

## Code Availability

The software utilized in the analysis of the data during the current study is not publicly available as it is property of Quantum-Si, Inc. Please contact the corresponding author, M.D.D., if you are interested in accessing the data.

## Material Availability

All materials are commercially available through Quantum-Si, Inc.

## Supplementary Information

### Semiconductor device operation and bin ratio calculation

Experiments were performed on pre-production semiconductor chips with 296K active wells, taking into account some loss to flow cell occlusion of the sensor array. A dual chamber flow cell allows for two independent samples to be sequenced in parallel, each utilizing 148K active wells. Initial production devices have 2M active wells, scaling to tens of millions of active wells using standard CMOS processing for the first product line. Pulsed 532 nm excitation light from a 67 MHz mode-locked laser is coupled into a grating coupler at the edge of the semiconductor chip. A network of optical waveguides divides the excitation light and routes it to the sensor array to illuminate each reaction chamber. Each CMOS pixel contains a single light-sensitive photodiode with two high-speed global shutters (a reject gate and a collect gate) that discard and collect photoelectrons. Control waveforms are applied to the collect and reject gates synchronously with the incident pulsed light source (Extended Data Fig. 1b). Approximately 1 ns before the excitation pulse, the reject gate is charged to >3 volts and the collect gate is discharged below 1 volt. Scattered 532 nm excitation photons generate photoelectrons in the photodiode. The photoelectrons are quickly transferred to a high voltage drain by built-in potential fields within the photodiode and the reject gate potential. Between 1 and 3 ns after excitation, the collect gate is charged to > 3 volts, the reject gate is discharged to < 1 volt. Photoelectrons generated from emitted photons that arrive in the photodiode after the collect gate is opened are transferred to a storage node within each pixel. Photoelectrons are accumulated for 7.5 to 30 ms, configurable, within each pixel across approximately 500,000 to 2,000,000 laser pulses (Extended Data Fig. 1b). The accumulated charge in the storage node is measured with the standard transfer gate, floating diffusion, source follower, row select, and on-chip analog-to-digital converters common to all CMOS image sensors, enabling scaling to large array sizes with small pixels. Fluorescence lifetime information is obtained by alternating the timing of the collect and reject gate waveforms between subsequent measurements. In the first measurement, only emission photoelectrons that arrive >3ns after the excitation pulse (bin 0) are collected. In the second measurement, emission photoelectrons that arrive >1ns after the excitation pulse (bin 1) are collected. The ratio of these two measurements (bin ratio) provides an estimate of the fluorescence lifetime (Extended Data Fig. 1c). Signal measured from the pixel as the phase relationship between the excitation source and the gate waveforms is adjusted throughout the entire excitation cycle demonstrates the pixel transitioning from 100% collection of photons during the collect phase to extinction of greater than 99.99% of photons during the rejection phase in less than 1ns (Extended Data Fig. 1d). We have demonstrated the ability to differentiate multiple dyes based on bin ratio alone (Extended Data Fig. 1e).

### Peptide synthesis and labeling

Peptides were synthesized on Rink Amide Resin on a PurePrep Chorus Solid-phase peptide synthesizer (Gyros Protein Technology) using Standard Fmoc chemistry. All synthetic peptides contained C-terminal Fmoc-azidolysine. The resin was deprotected in a mixture of TFA/TIPS/H2O (2.5%/2.5%/95%) at room temperature for 1.5 h. The deprotection mixture was concentrated under an argon stream. The peptides were precipitated from cold diethyl ether, resuspended in 1:1 water-acetonitrile, and purified on reverse phase HPLC (X-bridge C18, Waters) with a gradient of 10-70% acetonitrile (0.05% TFA) over 20 min. The residue was dried under high vacuum to generate white pellets. Into a solution of DBCO-DNA-biotin (2 nmol in 100 uL PBS) was added the peptide stock solution (4 uL, 5 mM) at room temperature. The reaction progress was monitored on LC-MS (Thermo UltiMate 3000 Executive Plus). After the reaction was completed, the mixture was conjugated to an excess of streptavidin. The peptide-DNA-streptavidin complex was purified on an ion exchange HPLC (DNAPac 200, Thermo). Gradient, buffer A, 20 mM sodium phosphate buffer, pH 8.5, buffer B, 1 M NaBr, 20 mM sodium phosphate buffer, pH 8.5, 20-60% B over 15 min. The purified complex was buffer-exchanged to a solution containing 50 mM MOPS (pH 8.0) and 60 mM potassium acetate on a 30K MWCO spin filter before use. The peptide containing fully oxidized methionine was prepared by mixing 3% hydrogen peroxide with the methionine peptide in 1:1 water-methanol at room temperature for 20 minutes. The product was immediately purified on a reverse-phase HPLC using the same peptide purification method described above, the purity was verified by reverse-phase HPLC (Thermo UltiMate 3000) on an analytical column (Zorbax SB-Aq, 5 μm, 4.6 × 250 mm), and the correct mass of the oxidized product was verified by LC-MS (Agilent LC-MSD-iQ, positive mode).

### Protein digestion and labeling

GLP-1 (7-37) and ubiquitin (1-76) recombinant proteins were purchased from RnD Systems as lyophilized powder. Each protein was reconstituted in 100 mM HEPES, pH 8.0 (20% acetonitrile) to a final concentration of 200 μM. When necessary, cysteines were reduced and alkylated using TCEP (2 mM) and iodoacetamide (10 mM). GLP-1 was digested using 1 μg of Trypsin (LCMS grade, Pierce) at 37 °C overnight. Ubiquitin was digested using 1 μg of LysC (LCMS grade, Pierce) and 1 μg of rAspN (LCMS grade, Promega). After protease digestion, pH of peptide mixtures was adjusted to pH 10.5 using potassium carbonate (57 mM), and lysines were converted to azidolysines using imidazole-1-sulfonyl azide (ISA, 2 mM) and copper sulfate catalyst (0.5 mM). ISA was quenched using polyurethane beads bearing an amine functionality (Oligo Factory). The mixture was then filtered and adjusted to pH 7-8 using 1 M acetic acid. The solution was diluted in 50% (v/v) of 10 mM MOPS, 10 mM KOAc, pH 7.5, added to DNA-streptavidin-DBCO complex, and incubated at 37 °C for 12-16 h. When required, the detergent Cetrimonium bromide was added to the reaction at a final concentration of 0.25 mM.

### Recognizer purification, labeling, and characterization

Expression vectors (with pET30 a+ backbone) for recognizers and Biotin ligase were co-transformed into BL21(DE3) chemically competent *E. coli* cells. The transformed cells were plated on Luria agar plates containing carbenicillin (50 μg/mL) and kanamycin (25 μg/mL) and incubated overnight at 37 °C to obtain single colonies. The starter liquid cultures inoculated with colonies were grown in Luria broth with ampicillin (50 μg/mL) and kanamycin (25 μg/mL) and inoculated into large cultures at a starting optical density (OD600) of ^~^0.01. The expression cultures were incubated at 37 °C at 230 rpm until OD600 approached ^~^0.7. The cultures were then induced with 0.4 mM IPTG. The expressed recognizer was biotinylated in vivo by adding 8 mM biotin at the same time as IPTG. Cells were harvested after ^~^12 hrs of expression by centrifugation at 10,000 *g* at 4 °C, and the cell pellets were washed with 1x PBS buffer pH 7.4. The cells were resuspended in Bug buster HT (Thermo Fisher Scientific) and incubated at room temperature for 30 mins on a magnetic stirrer. The cell suspension was then diluted with equal volume of 2x lysis buffer (100 mM Tris-HCl pH 7.5, 10% glycerol, 0.5 M NaCl) and incubated at room temperature for 30 mins on a magnetic stirrer. The lysate was centrifuged at 21,000 *g* at 4 °C to remove cell debris. Supernatant was collected and loaded on a Nickel NTA resin (Cytiva) affinity column pre-equilibrated with Buffer A (50 mM Tris-HCl pH 7.5, 10% glycerol, 0.5 M NaCl) on an AKTA Pure (Cytiva) system. The column was washed with at least ten column volumes of the buffer containing 10 mM imidazole. Elution was performed using a 10-300 mM imidazole gradient. Eluted fractions were dialyzed in a 10 kDa cassette against the dialysis buffer (50 mM Tris-HCl pH 7.5, 0.2 M NaCl, 50% glycerol) at 4 °C overnight.

For labeling of the recognizers, equal volumes of recognizer and DNA-Dye-Streptavidin complex were mixed at 5:1 (recognizer:DNA-dye-SV) molar ratio. The mixture was incubated on ice for 30 m and dialyzed overnight against SEC buffer (25 mM HEPES pH 8.0, 150 mM KCl). The recognizer-dye conjugate was harvested from the dialysis and centrifuged at 10,000 *g* at 4 °C. Supernatant was collected and concentrated using 10 kDa cut off concentrators. The concentrated conjugate was purified on an Agilent 1260 Infinity HPLC system using a size exclusion column (BioSEC-3 300 Å, 3 μm).

Binding affinity was measured by polarization using a labeled peptide. The polarization response and total intensity measurements were carried out at 20 °C on a microplate fluorometer with 480 nm excitation and 530 nm emission. The interaction of recognizer with labeled peptide containing a target N-terminal residue (XAKLDEESILKQK-FITC) was performed in PBS buffer at pH 7.4 and readings were collected after 30 min. Multiple analyses were performed at increasing recognizer concentration at a fixed concentration of a target peptide to obtain a titration curve. An equilibrium polarization response at each concentration was plotted and fit to calculate the *K*_D_.

The off-rate (*k*_off_) of PS610 was measured for various peptides using a stopped flow instrument. Labeled peptide (50 nM) was mixed with PS610 in PBS buffer pH 7.4 with 0.01% Tween-20 and incubated at 30 °C. After 30 min of incubation, the recognizer:peptide complex was rapidly mixed with 10-20 fold molar excess of unlabeled trap peptide and the reaction was followed in real-time by measuring the fluorescence intensity. At least three time-course traces were averaged and fit to an exponential equation.

### Aminopeptidase purification

Expression vectors (with pET30 a+ backbone) for aminopeptidases PhTET2 and PhTET3 were transformed into BL21(DE3) chemically competent *E. coli* cells. The transformed cells were plated on Luria agar plates containing kanamycin (25 μg/mL) and incubated overnight at 37 °C to obtain single colonies. The starter liquid cultures inoculated with colonies were grown in Luria broth (LB) with kanamycin (25 μg/mL) and inoculated into large cultures at a starting optical density (OD600) of ^~^0.01. The expression cultures were incubated at 37 °C at 230 rpm until OD600 approached ^~^0.7. The cultures were then induced with 0.4 mM IPTG. The expressed aminopeptidase was purified as described above for recognizers. For conditioning, the aminopeptidase protein was dialyzed against 50 mM MOPS pH 8.0/ 60 mM potassium acetate and then exposed to cobalt acetate at a final concentration of 400μM for 1-1.5 h at 65 °C to form the active dodecamer complex. The conditioned aminopeptidase preparation was dialyzed further against 50mM MOPS pH 8.0/ 60 mM potassium acetate, aliquoted, and flash frozen.

### Peptide loading, recognition, and dynamic sequencing

The semiconductor chip was placed in the sequencing device and a chip check was performed to test electronic circuit function and to optimize laser coupling alignment. The chip was then removed from the device socket and the chip was washed twice with 50 μL of 70% isopropanol, followed by four washes with 30 μL of wash buffer (50 mM MOPS pH 8.0, 60 mM potassium acetate, 50 mM glucose, 20 mM magnesium acetate, and surfactant mix) through a flow cell attached to the chip. A second chip check was then performed. The laser was then blocked via an integrated software-controlled shutter, peptide complex was added to one half of the chip to a final concentration of 1-10 nM and mixed thoroughly, and the chip was incubated for 15 min. The chip was then washed six times with wash buffer, followed by addition of an imaging solution (wash buffer with 5 mM Trolox and an oxygen scavenging system). The laser was unblocked and the occupancy percentage (target 10-30%, Poisson distributed) was recorded by acquiring a photobleaching signal from a fluorophore attached to the peptide complex during 5 min of laser illumination. For NAA recognition-only assays, after peptide loading, labeled recognizer was added to a final concentration of 50 nM PS610, 100 nM PS691, or 250 nM PS961 (as indicated according to the experiment), and data was recorded for 10 hours. For dynamic sequencing assays, after peptide loading, a mixture of labeled recognizers was added to obtain final concentrations of 50 nM PS610, 100 nM PS691, and 250 nM PS961. Data was recorded for 15 min. The laser was then blocked briefly and aminopeptidases were added to the sequencing reaction via the flow cell and mixed thoroughly (final concentration 2-8 μM PhTET2 and/or 20-80 μM PhTET3, as indicated according to the experiment). The laser was then unblocked, and data was recorded for 10 hours. For all runs, 30 μL of mineral oil was added to fluid reservoirs at each port of the flow cell to prevent evaporation during the run.

### Signal processing and trace segmentation

The measured signal on-chip comprises various noise components, the most dominant one being due to fluorescent emissions from diffusing recognizers in the reaction chamber. The pulse caller algorithm for a given reaction chamber starts by estimating the statistical properties of this background noise component. Once an estimate within certain error bounds has been established, the algorithm works in an online fashion observing new frames of data as they are generated. At each point in time, the algorithm maintains state indicating whether the signal is due to the background component only or a pulse from a recognizer-NAA interaction is being observed. The state transition from background to pulse is triggered using an edge detection test where the shift in signal is expected to be significant with respect to the background component’s statistical distribution. The state transition from pulse to background is triggered when a small window of the most recent frames of the signal appears to conform to the background component’s distribution again. The algorithm maintains an updated model of the background component as new background frames are observed. This provides robustness against drift in the signal intensity together with a feedback control loop that maintains a stable optical coupling of the laser into the chip based on any such detected drift. As detected pulses can be due to true recognizer-to-dipeptide interaction events as well as other occasional transient noise spikes, a downstream filter layer is employed to test the significance of pulse events based on their duration, intensity, and noise patterns within the context of the full timeline of the run and the entire dataset of reaction chambers.

Initial regions are determined by performing a sliding window calculation of pulse rate along the time dimension of a series of pulses. Regions with a mean pulse rate > 1 pulse/min are then subdivided according to a greedy bisection approach in which all potential splits are assessed by a Mann-Whitney U Test to find statistically significant deviations between pulse properties (intensity, time bin ratio, pulse duration, and interpulse duration) in the pulses on the left and right of the split. To define RSs, the split point with the lowest p-value is used to sub-divide the region and the process continues until no regions remain with a candidate split point with p-value < 10^-5^.

### Recognition segment classification

RS classification for reactions containing single synthetic peptides was performed using an unsupervised clustering algorithm. A subset of RSs including those with mean signal-to-noise ratio of their constituent pulses of ≥3 were used to pre-train a Gaussian mixture model (GMM) to identify approximate centroids for each of N classes of recognition, where N equals the number of expected recognizable peptide states with F, Y, W, L, I, V, or R at the N-terminus. Identified clusters were assigned to recognizable peptide states by matching the predominant order of cluster sequences observed to the expected amino acid sequence and by using prior knowledge of dye properties to identify the recognizers active during each RS. Subsequent rounds of GMM fitting were performed on all RSs matching the expected order of these events to refine the GMM model until no further sequences appeared in the expected order. The final model was then applied to all RSs in a given reaction.

RS classification for reactions containing library prepared peptides and mixes of peptides was performed using a random forest classifier that was pre-trained on annotated RS pulse features from prior synthetic peptide experiments. Unless otherwise noted, figures and statistics produced from classified RSs are derived from reaction chambers containing the expected sequence of RSs.

### Molecular dynamics and binding energy calculation

Homology models of PS961 complexed to peptide were generated using an internal crystal structure, mutations were applied and optimized using protCAD^22^ prior to molecular dynamics. AMBER20^23^ implicit solvent molecular dynamics simulations using the generalized Born^24^ solvation potential were performed using the ff19SB^25^ force field with no atomic distance cutoff. Minimization was performed using steepest descent, followed by conjugate gradient minimization. The system was thermalized from 0 to 300K using Langevin dynamics and a collision frequency of 3 ps^-1^. Molecular dynamics simulations of the equilibrated recognizer-peptide complex, free recognizer and free peptide were independently run for 5 nanoseconds at 300 K to perform the binding energy calculation using MMPBSA^26^. Where 125 frames, each containing 10,000 2 femtosecond steps, were used for the calculation from the three simulations. Binding energy and the decomposition of all residues contributing to the binding energy was computed in 0.15 M salt concentration.

### Protein identification

Human proteins (UniProtKB/Swiss-Prot H. sapiens proteome UP000005640) were digested in silico at cleavage sites for AspN and LysC to obtain a database of all peptide fragments with C-terminal lysine. The mature 76-amino acid ubiquitin sequence was used to represent all proteins encoding ubiquitin precursors. Candidate matches to the kinetic signature for DQQRLIFAGK peptide obtained from ubiquitin sequencing were identified by searching for a pattern supported by kinetic information in the kinetic signature: the RLIF motif preceded by at least 2 non-proline residues, followed by at least one residue, and not containing the recognizable amino acids F, Y, W, L, I, V, or R, in the sequence flanking the RLIF motif.

## Extended Data Figures

**Extended Data Fig. 1.**
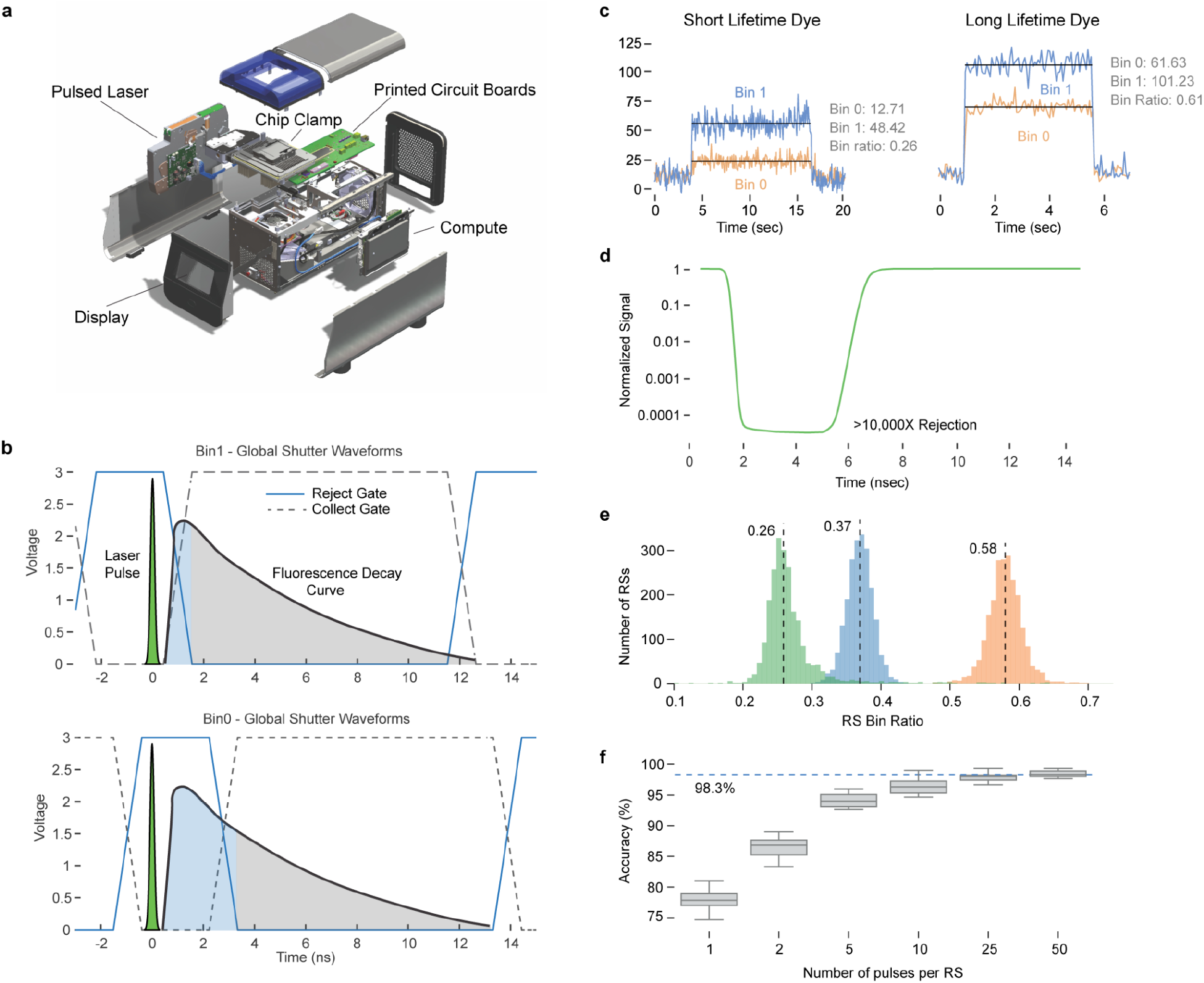
Chip operation. **a**, Exploded view of the compact benchtop instrument designed to support the custom semiconductor chip and protein sequencing assay. **b**, The chip achieves electronic rejection by discarding photoelectrons from the pulsed laser before shifting to collect fluorescence photoelectrons from bound NAA recognizers. The timing of the rejection and collection windows cycles between two modes (Bin 1 and Bin 0, example waveforms shown) in alternate frames to provide a bin ratio estimate of the fluorescence lifetime of the dye. **c**, Example pulses for dyes with short and long fluorescence lifetime, illustrating the difference in signal collection in Bin 0 and Bin 1. **d**, The chip achieves >10,000-fold attenuation of incident laser light within 1 ns from initiation of a rejection mode. **e**, Distributions of mean RS bin ratio collected for three dyes with different fluorescence lifetime. **f**, Dye channel identification accuracy increases with the number of pulses captured per RS.

**Extended Data Fig. 2.**
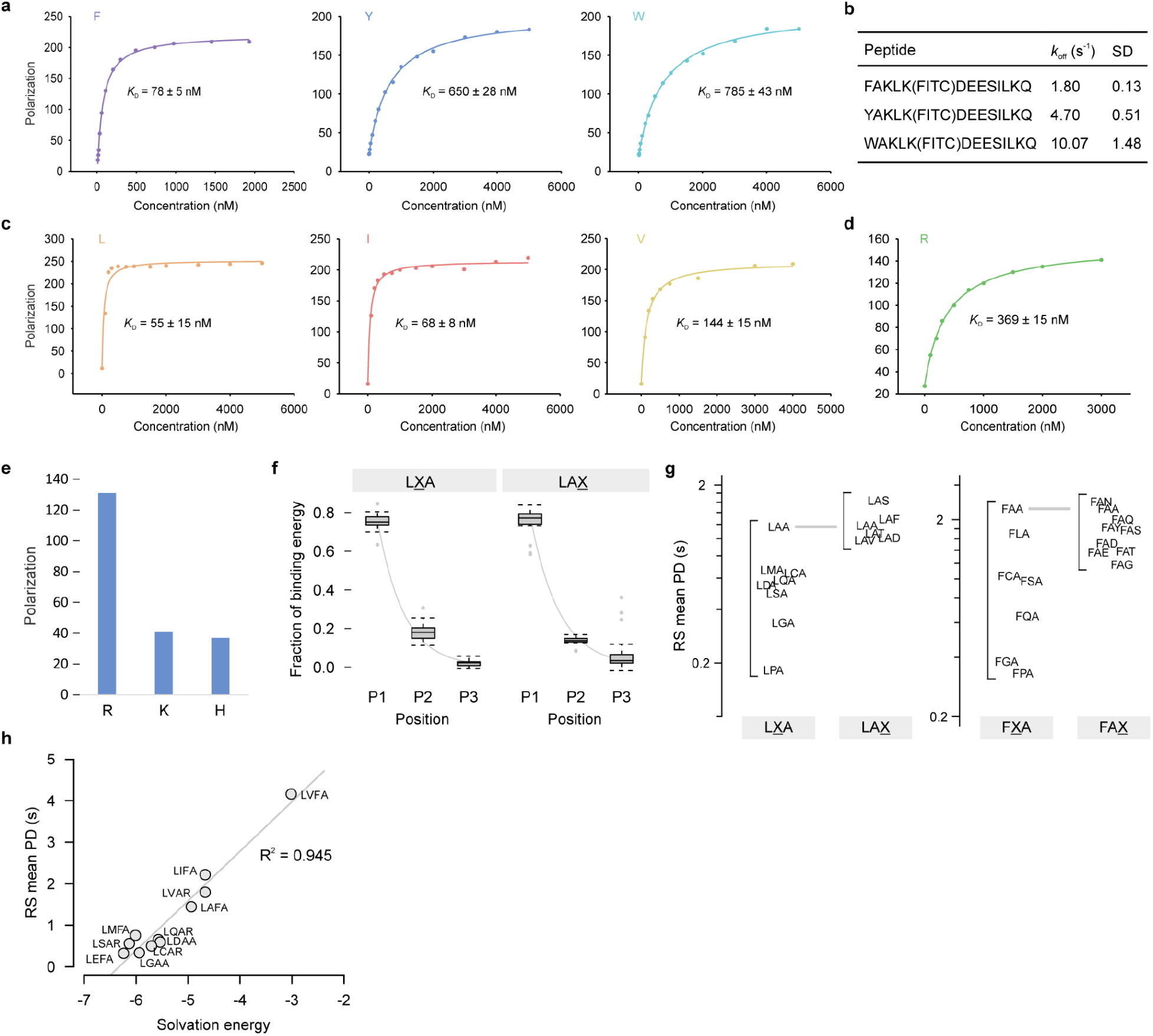
Recognizer properties. **a-e**, Recognizer kinetic characterization using polarization assays (Methods). **a,b**, Affinity (*K*_D_) and off-rate (*k*_off_) of PS610 for peptides with N-terminal phenylalanine, tyrosine, and tryptophan. **c**, Affinity of PS961 for peptides with N-terminal leucine, isoleucine, and valine. **d,e**, Affinity of PS691 for a peptide with N-terminal arginine (**d**) and single-point polarization data measured for peptides with N-terminal arginine, lysine, and histidine (**e**). **f**, Binding energy was calculated using a computational model (Methods) for peptides of initial sequence LXA and LAX, where X = all 20 amino acids. Boxplots show the fraction of total binding energy contributed by the amino acid at P1, P2, and P3, with an exponentially decreasing trend from P1 to P3 (R^2^ > 0.97). **g**, RS mean PD determined in single-molecule assays for LXA and LAX peptides using PS961 and for FXA and FAX peptides using PS610. **h**, The non-polar solvation energy term from the computational binding model with PS961 exhibits high correlation with actual RS mean PD values observed in single-molecule assays with peptides containing N-terminal leucine and varying widely at P2.

**Extended Data Fig. 3.**
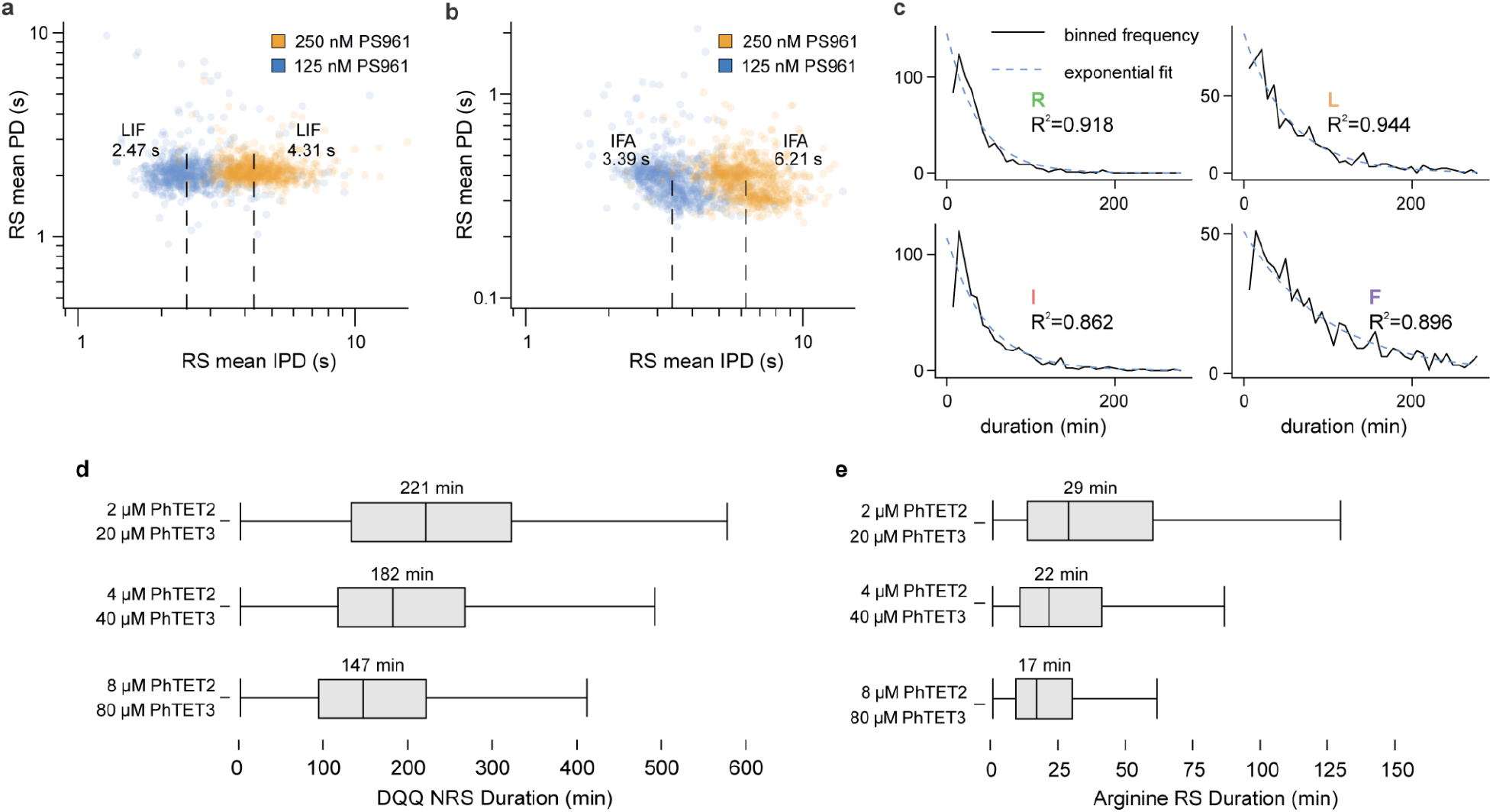
Binding and cleavage rates. **a,b**, RS mean IPD for PS961 binding to LIF and IFA in dynamic sequencing assays at a concentration of 125 nM or 250 nM. Median IPD values are indicated. **c**, Single exponential decay curves fit to the RS duration distributions for arginine, leucine, isoleucine, and phenylalanine acquired from dynamic sequencing of the synthetic peptide DQQRLIFAG. **d,e**, Increasing the aminopeptidase concentrations in dynamic sequencing runs of the synthetic peptide DQQRLIFAG resulted in decreased NRS (left) and RS (right) durations. Median RS duration values are indicated.

**Extended Data Fig. 4.**
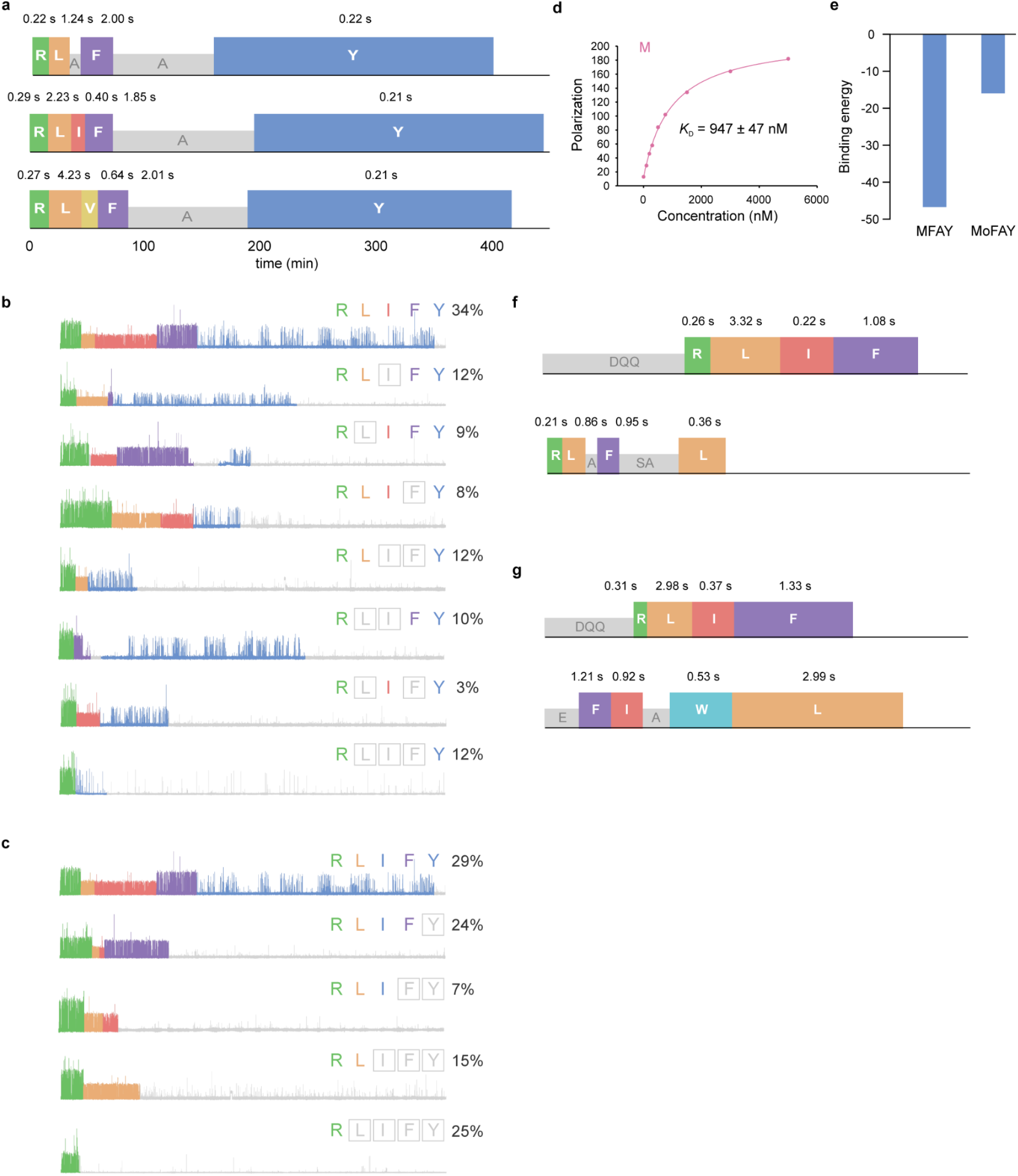
Kinetic signatures from single amino acid changes and PTMs. **a**, Kinetic signature plots for three peptides: RLAFAYPDDD (top), RLIFAYPDDD (middle), and RLVFAYPDDD (bottom). **b-c**, Incomplete RS information observed in dynamic sequencing of RLIFAYPDDD peptide. **b**, Percentage of reads and example traces of each type of observed deletion of one or more RSs in traces beginning with arginine and ending with tyrosine recognition. **c**, Percentage of reads and example traces of each type of observed truncation of one or more RSs in traces beginning with arginine. **d**, Affinity of PS961 for a peptide with N-terminal methionine measured using a polarization assay (Methods). **e**, Binding energy prediction for peptides with N-terminal methionine and methionine sulfoxide (Mo) from computational modeling with PS961 (Methods). **f**, Kinetic signature plots for DQQRLIFAG and RLAFSALGAADDD peptides mixed and run on the same chip. **g**, Kinetic signature plots for DQQRLIFAGK and EFIAWLVK peptides obtained from digestion of recombinant human ubiquitin and GLP-1.

